# Transcriptomics, Regulatory Syntax, and Enhancer Identification in Heterogenous Populations of Mesoderm-Induced ESCs at Single-Cell Resolution

**DOI:** 10.1101/2022.03.29.486247

**Authors:** Mamduh Khateb, Jelena Perovanovic, Kyung Dae Ko, Kan Jiang, Xuesong Feng, Natalia Acevedo-Luna, Jérome Chal, Veronica Ciuffoli, Pavol Genzor, James Simone, Astrid D. Haase, Olivier Pourquié, Stefania Dell’Orso, Vittorio Sartorelli

## Abstract

ESCs can adopt lineage-specific gene expression programs by stepwise exposure to defined factors, resulting in the generation of functional cell types. Bulk and single cell-based assays were employed to catalogue gene expression, histone modifications, chromatin conformation, and accessibility transitions in ESC populations and individual cells acquiring a presomitic mesoderm fate and undergoing further lineage specification. These assays identified *cis*-regulatory regions and transcription factors presiding gene expression programs occurring at defined ESC transitions and revealed the presence of heterogeneous cell populations within discrete ESC developmental stages. The datasets were employed to identify previously unappreciated genomic elements directing the initial activation of Pax7 and myogenic and neurogenic gene expression programs. This study provides a resource for the discovery of genomic and transcriptional features of pluripotent, mesoderm-induced ESCs, and ESCs-derived cell lineages.

## INTRODUCTION

Embryonic stem cells (ESCs) self-renew indefinitely, retain pluripotency, and differentiate into all adult cell types without changes in their genetic information. DNA and histone chemical modifications as well as chromatin accessibility and architecture collectively ensure genome stability, correct propagation of genetic information, and proper interpretation of the genome (Allis and Jenuwein, 2016). Enhancers play a key role in these processes by orchestrating cell type-specific gene expression programs (Visel et al., 2009) (Catarino and Stark, 2018). Muscle stem cells (MuSCs), which are required for muscle growth and regeneration (Brack and Rando, 2012) (Yin et al., 2013) (Tierney and Sacco, 2016) (Evano and Tajbakhsh, 2018) (Fuchs and Blau, 2020) (Relaix et al., 2021), can be generated from mouse or human pluripotent stem cells by either treatment with defined molecules or forced expression of the transcription factor Pax7 (Darabi et al., 2012) (Chal et al., 2015) (Chal et al., 2018) (Al Tanoury et al., 2020) (Xi et al., 2020) (Magli and Perlingeiro, 2017). Using a protocol mimicking key signaling events occurring in the mouse embryo (Chal *et al*., 2015), we induced ESCs differentiation and characterized their transcriptome and epigenome at specific developmental junctures. Integration of these data uncovered chromatin dynamics underlying activation of cell fate programs in ESC-derived precursors confirmed to occur in somites of mouse developing embryos. Bulk and single cell analysis revealed distinct molecular syntax utilized by regulatory regions to control expression of individual genes within a common gene regulatory network. We leveraged these datasets and, by combining chromatin accessibility, *in situ* Hi-C chromatin conformation, genome editing, and reporter assays have identified regulatory regions directing initial Pax7 expression and activation of the myogenic and neurogenic programs.

## RESULTS

### Instructing ESCs

Mouse ESCs in which the green fluorescent protein GFP coding region has been placed under the control of endogenous Pax3 regulatory sequences (ESCs Pax3-GFP) (Chal *et al*., 2015) were cultured in conditions maintaining pluripotency (feeder-free conditions, LIF+2i inhibitors, naïve ESCs) (Ying et al., 2008). BMP4 is essential for embryogenesis, predominantly for mesoderm formation (Dale et al., 1992), and drives commitment to differentiation of human ESCs (Gunne-Braden et al., 2020). We induced initial ESCs differentiation employing a medium supplemented with Bmp4 protein (10ng/ml) and 1% KSR for 48 hr (instructed ESCs) (see Experimental Procedures) (Figure 1A and Supplemental Figure S1A). RNA-seq analysis (1.5-fold change, adjusted p<0.05) revealed 2,022 upregulated and 1,288 downregulated genes in instructed ESCs (Figure 1B, Supplemental Table S1). Gene ontology of upregulated transcripts returned terms related to cholesterol biosynthesis, cell morphogenesis, steroid biosynthesis, actin cytoskeleton organization, and Rho GTPase cycle (Supplemental Figure S1B, Supplemental Table S1). Down-regulated transcripts in instructed ESCs were enriched for terms related to lysosome, autophagy, and pluripotency (Supplemental Figure S1B, Supplemental Table S1). Transcription factors Otx2 and Pou3f1 (Oct6), fibroblast growth factors Fgf5 and Fgf15, DNA methyltransferases Dnmt3a, Dnmt3b, and secreted factor Wnt8a, which mark an intermediate state (formative pluripotency) of ESCs transitioning from naïve to primed pluripotency *in vitro* and *in vivo* (Buecker et al., 2014) (Boroviak et al., 2015) (Acampora et al., 2016) (Kinoshita et al., 2021) (Wang et al., 2021) (Pera and Rossant, 2021) were upregulated in instructed ESCs (Figure 1C,E,F). Moreover, a core of transcripts enriched in human pluripotent stem cells skewed toward a naïve-to-primed intermediate state (Cornacchia et al., 2019) was also increased in instructed ESCs (UP-lipid_signaling, Supplemental Table S1). Instead, pluripotency factors Nanog, Esrrb, Klf4, Prdm14, Tbx3, and Zfp42, downregulated in ESCs with epiblast-like primed pluripotency (Buecker *et al*., 2014), were reduced (Figure 1D,E,F). Altogether, these data indicate that Bmp4-instructed ESCs share gene expression profiles with ESC formative-intermediate states.

**Figure 1.**
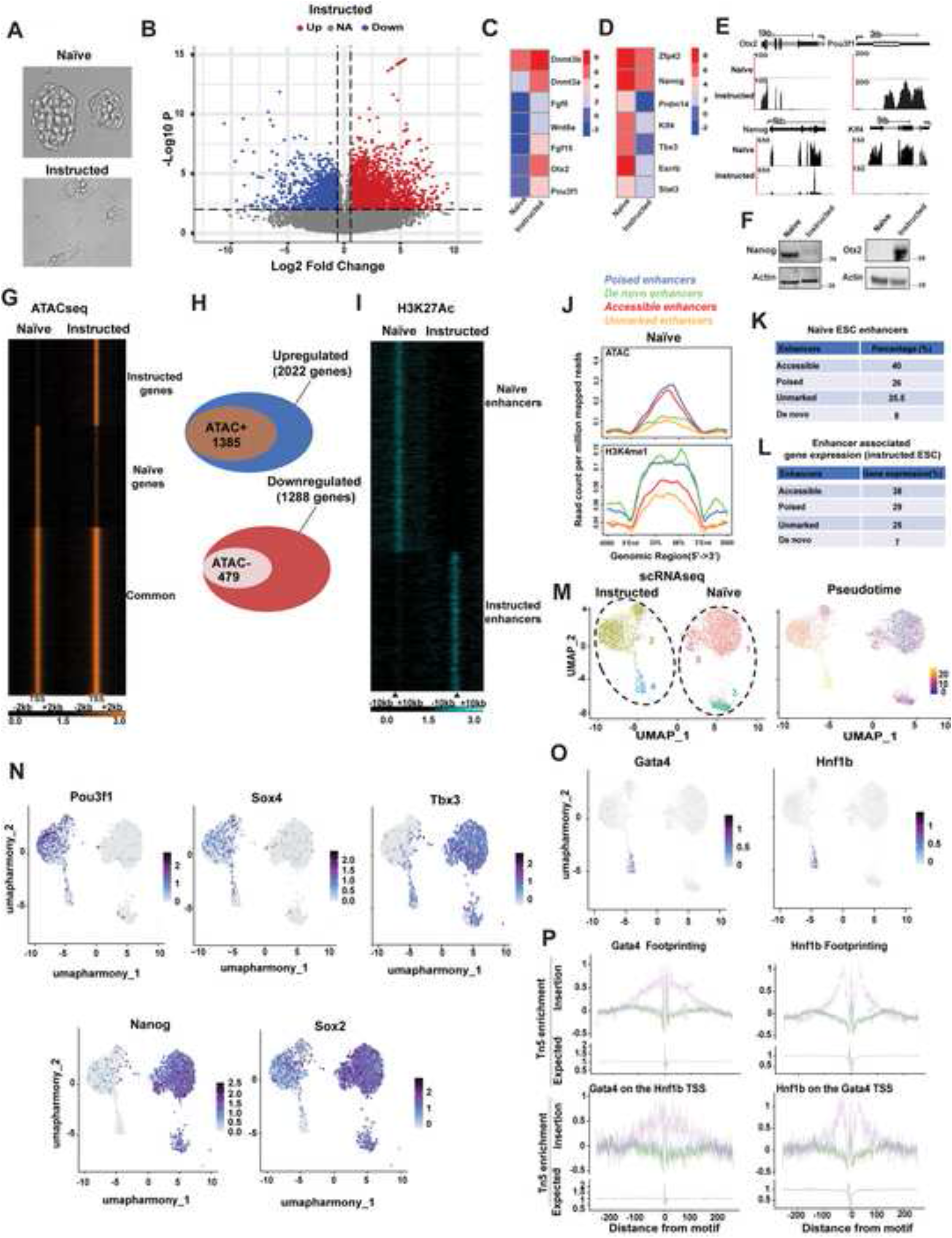
Bulk and Single-Cell Analyses of Transcriptomes, Enhancers, and Chromatin Accessibility in Naïve and Bmp4-Treated ESCs. (A) Morphology of ESCs cultured 2i+LIF conditions (Naïve) or exposed to Bmp4 in serum-free medium for 48 hr (Instructed). (B) Transcriptome changes of instructed ESCs. Each point represents RPKM values obtained for a given transcript in RNA-seq analysis from instructed ESCS. Abscissa represents magnitude of Log2 fold change and ordinate indicates statistical significance (-Log10 P-value). (C,D) Heatmaps representing expression of selected transcripts in naïve and instructed ESCs. (E) RNA-seq tracks of Otx2, Pou3f1, Nanog, and Klf4 in naïve and instructed ESCs. (F) Immunoblot of Nanog or Otx2 protein in naïve and instructed ESCs. (G) Heatmap representing ATAC-seq signal intensity in naïve and instructed ESCs. Signal is centered −2Kb/+2Kb from the transcriptional start site (TSS). (H) Venn diagram intersecting TSS of upregulated genes acquiring ATAC-seq signal (top panel) and TSS of downregulated genes losing ATAC-seq signal in instructed ESCs (bottom panel). (I) H3K27ac heatmap in naïve and instructed ESCs. Interval is −10Kb/+10Kb from the center of the peak signal. (J) Averaged normalized tag intensities of ATAC-seq (top panel) or H3K4me1 (bottom panel) signal at genomic regions in naïve ESCs acquiring H3K27ac in instructed ESCs. (K) Percentage of genomic regions with distinct ATAC-seq and H3K4me1 enrichment in naïve ESCs acquiring H3K27ac signal in instructed ESCs. (L) Percentage of upregulated genes in instructed ESCs associated with genomic regions with distinct ATAC-seq and H3K4me1 enrichment in naïve ESCs. (M) scRNA-seq UMAP of naïve and instructed ESCs (left panel). Pseudotime ordering of naïve and instructed ESCs clusters (right panel). (N) Expression of Pou3f1, Sox4, Tbx3, Nanog, and Sox2 transcripts in naïve and instructed ESCs clusters. (O) Expression of Gata4 and Hnfb1 in naïve and instructed ESCs clusters. (P) Overall footprinting of Gata4 and Hnf1 in instructed ESCs (top panel), and Gata4 footprinting at Hnf1b TSS and Hnfb1 footprinting at Gata4 TSS, respectively (bottom panel).

### Identification of the Regulatory Landscape in Instructed ESCs

Chromatin accessibility to transcription factors and cofactors is essential for establishing and maintaining cellular identity (Klemm et al., 2019). Surveying chromatin accessibility of naïve and instructed ESCs by ATAC-seq revealed that 68% (1,385/2,022) of genes upregulated in instructed ESCs gained increased chromatin accessibility at their TSS (Figure 1G,H). Of the 1,288 down-regulated genes, 37% (479/1,288) displayed reduced chromatin accessibility (Figure 1G,H). Cell type- and cell state-specific gene expression requires coordination between promoters and enhancers (Field and Adelman, 2020). To identify active enhancers, we conducted H3K4me1 and H3K27ac ChIP-seq (Figure 1I) and integrated the resulting datasets with ATAC-seq. Of the 1,482 instructed ESCs-specific enhancer regions (defined as H3K4me1^+^/H3K27ac^+^/ATAC^+^), 25% were associated with increased transcription of the nearest gene (Supplemental Table S1 and S2). Albeit observed in a different experimental setting, these results agree with a study reporting that only a subset of genomic regions marked by H3K4me1 and H3K27ac in ESCs displayed enhancer activity (Barakat et al., 2018). Enhancers were activated with different modalities. In naïve ESCs, 40% (597/1,482) of accessible enhancers had low H3K4me1 and were ATAC^+^, 26% (386/1,482) of poised enhancers were H3K4me1^+^/ATAC^+^, 25.5% (379/1,482) of unmarked enhancers had the lowest H3K4me1^-^ and were ATAC^-^, and 8% (120/1,482) of *de novo* instructed ESCs enhancer*s* were H3K4me1^+^/ATAC^-^ (Figure 1J,K and Supplemental Table S2). All 1,482 regions acquired H3K27ac in instructed ESCs (Supplemental Figure S1C). Accessible enhancers were associated with 38%, poised enhancers with 29%, unmarked enhancers with 25%, and *de novo* enhancers with 7% of genes activated in instructed ESCs, respectively (Figure 1L). Here below, we provide selected examples of enhancer activation modalities. Enhancer and promoter regions of up-regulated genes Pou3f1 (Oct6) and Wnt8a gained chromatin accessibility and H3K27ac (Supplemental Figure S1D). Intronic enhancer regions of upregulated gene Pbx1 and Pitx2 genes were already accessible in naïve conditions and acquired H3K27ac only upon ESCs instruction. Selected pluripotent genes were downregulated in instructed ESCs (Figure 1D). H3K27ac decreased while chromatin accessibility was preserved at enhancers of Esrrb gene. H3K27ac and chromatin accessibility were both reduced at upstream regulatory regions of Nanog gene. In another instance, pluripotent Tbx3 gene was downregulated, enhancer and promoter regions showed reduced H3K27ac and decreased chromatin accessibility. Thus, concordantly transcribed genes can be differently regulated. While frequently consonant with gene expression, chromatin accessibility and H3K27ac can be temporally disjointed, with chromatin accessibility often preceding transcription.

### Single Cell Analysis of Naïve and Instructed ESCs

Because of the stochastic nature of the pluripotency network, ESCs are transcriptionally heterogeneous even when exposed to a uniform culture environment (Torres-Padilla and Chambers, 2014). Establishing whether extracellular signals monotonously induce gene expression changes across a whole cell population or elicit pliable transcriptional responses in individual cells requires single cell approaches. Uniform manifold approximation and projection (UMAP) was employed to construct a high dimensional graph representation of scRNA-seq datasets obtained from naïve and instructed ESCs, respectively. Integration of the scRNA-seq datasets resulted in the identification of spatially distinct clusters that could be assigned to either naïve or instructed ESCs and placed on a pseudotemporal map (Figure 1M). The main naïve ESCs cluster (Figure 1M, cluster 1) expressed high levels of pluripotency genes, oxidative phosphorylation, and glycolytic transcripts, in line with the observation that pluripotent ESCs rely on both metabolic pathways for energy and metabolites production (Zhou et al., 2012) (Ryall et al., 2015) (Supplemental Table S2). Autophagy maintains ESCs stemness and Ulk1 (Atg1) is essential for this process (Gong et al., 2018). Expression of autophagy genes Ulk1 and Gabarapl2 (Atg8) was enriched in naïve ESC clusters (Supplemental Figure S1E). Two smaller additional clusters observed in naïve ESCs (Figure 1M, clusters 3 and 5) were also enriched for pluripotency, ribosomal, and translational transcripts (Supplemental Table S2). Instructed ESCs generated two clusters located at distinct pseudotemporal points. The main instructed ESC cluster (Figure 1M, cluster 2) occupied an intermediate pseudotemporal space between naïve ESCs and an additional instructed ESC cluster (cluster 4) and contained cells enriched for TFs Pou3f1 (Oct6), Sox4, Otx2, and Sall2 (Figure 1N and Supplemental Figure 1F). Pluripotency genes displayed a graded expression in the different clusters. While overall reduced compared to naïve ESCs, Nanog and Sox2 expression was still maintained in cluster 2 but drastically reduced in cluster 4 (Figure 1N). In contrast, Tbx3 displayed a bimodal expression in naïve cluster 1 and in instructed cluster 4, respectively (Figure 1N). This observation is consistent with a dual Tbx3 role in maintaining pluripotency and directing a cell-fate decision towards mesoendoderm (Weidgang et al., 2013). Instructed ESC clusters hosted cells sharing ∼ 28% of genes belonging to E8.5 and E9.5 neuromesodermal progenitor signatures (Gouti et al., 2017) (Supplemental Table S2). Gata transcription factors (TFs) can bind condensed chromatin and function as pioneer factors (Cirillo et al., 2002) (Sharma et al., 2020) and are co-expressed with the bookmarking TFs Hnf1 involved in the mitotic transmission of epigenetic information (Verdeguer et al., 2010). Expression of Gata4, Gata6, Foxa2, and Hnf1b, involved in mesoendoderm induction (Rojas et al., 2005) (Wamaitha et al., 2015) (Coffinier et al., 1999) (Costello et al., 2015) were enriched in cluster 4 (Figure 1O and Supplemental Figure S1H). scATAC-seq revealed that DNA binding motifs (DBMs) for Gata4, Gat6, Foxa2 and HNF1b were overrepresented and footprinted. Gata4 and Hnf1b reciprocally occupied their TSS. (Figure 1P and Supplemental Figure S1G,H). Thus, a single cell approach documented transitional states occurring in ESCs exposed to Bmp4 and identified TFs involved in these transitions.

### Single Cell Analysis of Pax3-Expressing Cells

Instructed ESCs were switched to culture conditions favoring acquisition of anterior presomitic mesoderm (aPSM) fate (R-spondin3 and the Bmp inhibitor LDN193189, RDL, see Experimental Procedures) (Supplemental Figure 2A) (Chal *et al*., 2015). After 96 hr in aPSM-promoting medium, ∼ 30-40% of ESCs Pax3-GFP became GFP^+^ and could be FACS-isolated (Figure 2A,B, aPSM cells). scRNA-seq analysis identified four cell clusters that were ordered on a pseudotime map (Figure 2C). Posterior PSM (pPSM) genes were not expressed (Figure 2D) while aPSM genes were highly expressed in the main cell clusters (clusters 0,1) (Figure 2E). Integration of scRNA-seq and scATAC-seq data, revealed that, while not transcribed, the chromatin of pPSM genes was accessible (Figure 2D). Since pPSM genes are expressed in ESCs cultured in aPSM-promoting medium for 48hr (Chal *et al*., 2015), chromatin accessibility at their promoter regions may indicate memory preservation of a previously active, now extinct, gene program. For instance, pPSM Wnt5a, Wnt5b, Hoxd11, and Hoxd13 genes were not expressed while their chromatin was accessible (Figure 2D). Fgf8, which activates Hoxd11 and Hoxd13 (Rodrigues et al., 2017), was hardly detected in aPSM cells (Supplemental Figure S2B) potentially explaining lack of Hoxd11-Hoxd13 activation despite their permissive chromatin. Two additional clusters were identified (Figure 2C). Cluster 3 was composed of cells expressing vessel-related genes vascular endothelial growth factor receptors Flt1, Flt4, and Kdr (Figure 2F and Supplemental Figure S2C). Cluster 2 hosted cells enriched for glycolytic genes and neuronal markers Pax6, Sox2, Sox11, and Nestin (Figure 2G,H, and Supplemental Figure S2D). Pax6, Sox2, and Nestin are highly expressed in neural stem cell precursors (NSCs) with Pax6 and Sox2 controlling NSCs identity and differentiation (Gotz et al., 1998) (Gomez-Lopez et al., 2011). Glycolysis prevents NSCs precocious differentiation (Lange et al., 2016) raising the possibility that activated glycolysis may prevent further differentiation of ESC-derived neurogenic cells to intermediate progenitors. Sox2 was expressed in both naïve ESCs and in aPSM cluster 2 (Figure 2 I). Sox2 is regulated by pluripotency TFs in ESCs (Young, 2011) and by Pax6 in neurogenic cells (Wen et al., 2008). Thus, we evaluated whether distinct Sox2 regulatory regions may be accessible in naïve ESCs and aPSM cells by scATAC-seq (Figure 2J,K). Enhancers located ∼ 100Kb from the Sox2 TSS (SRRs) (Zhou et al., 2014) were accessible in naïve ESCs (Figure 2K). Two other regions next to SRRs were also accessible in naïve ESCs (Figure 2K, red lines). In aPSM ESCs, all these elements became inaccessible while increased accessibility was observed at two regions located at ∼ 10Kb from the Sox2 TSS (Figure 2K). One of these regions, the N1 enhancer, directs Sox2 expression in the posterior neural plate (Takemoto et al., 2006). Another region (R1), located between N1 and SRRs enhancers, gained accessibility in aPSM cells (Figure 2K). Thus, besides expressing aPSM genes, ESCs prompted to acquire a PSM-like fate can also activate gene programs observed in endothelial and neurogenic precursors. Neurogenic precursors derived from ESCs recapitulate Sox2 enhancer regulation observed in the embryo’s neural plate.

**Figure 2.**
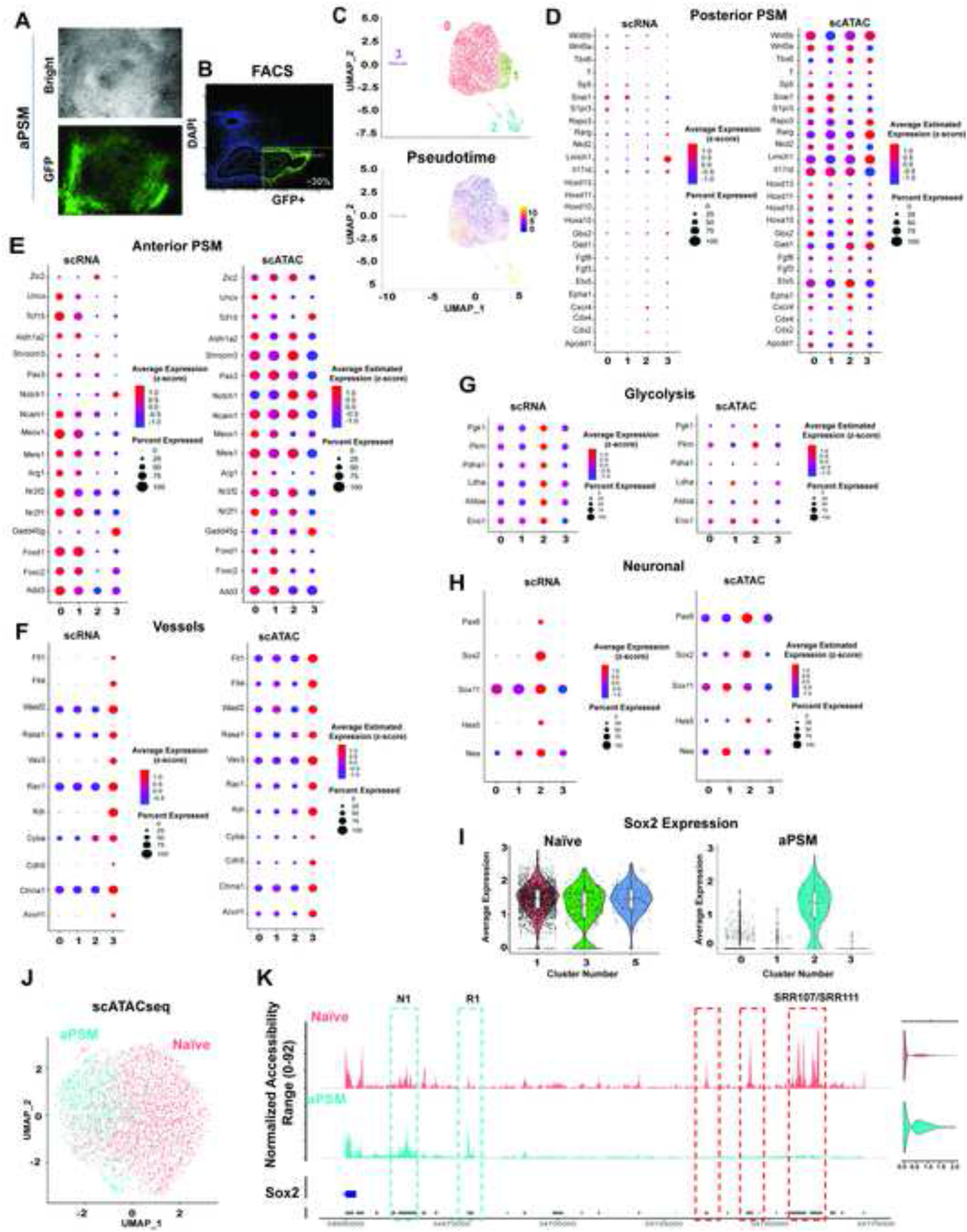
Single Cell Analysis of Transcriptome and Chromatin Accessibility of Pax3-GFP-Positive ESCs. (A) Bright-field and fluorescence microscopy of ESCs Pax3-GFP cultured in conditions favoring acquisition of anterior presomitic mesoderm (aPSM) fate. (B) FACS isolation of GFP^+^ ESCs Pax3-GFP. (C) scRNA-seq UMAP graph of GFP^+^ ESCs Pax3-GFP (top panel) and pseudotime ordering (bottom panel). (D) scRNA-seq and scATAC-seq Louvain dot plot of selected posterior PSM markers. (E) scRNA-seq and scATAC-seq Louvain dot plot of selected anterior PSM markers. (F) scRNA-seq and scATAC-seq Louvain dot plot of selected vessel endothelial markers. (G) scRNA-seq and scATAC-seq Louvain dot plot of selected glycolytic genes. (H) scRNA-seq and scATAC-seq Louvain dot plot of selected neuronal markers. (I) Violin plots representing Sox2 expression in naïve and GFP^+^ ESCs Pax3-GFP (aPSM). (J) scATAC-seq clustering of in naïve and GFP^+^ ESCs Pax3-GFP (aPSM). (K) scATAC-seq tracks of the Sox2 locus in naïve and GFP^+^ ESCs Pax3-GFP (aPSM). SRR107/SRR111 indicate Sox2 enhancers active in naïve ESCs. N1 denotes a Sox2 enhancer active in neurogenic cells. Next to N1, a genomic region (R1) with increased chromatin accessibility in aPSM cells. Violin plot quantification of Sox2 chromatin accessibility in naïve and aPSM cells.

### Identification of ESC-Derived Cell Lineages by Single Cell Omics

ESCs cultured in aPSM-promoting medium were switched to a medium supplemented with hepatocyte growth factor (HGF), insulin growth factor 1 (IGF-1), fibroblast growth factor 2 (FGF-2), LDN193189, and Rspo-3 (HIFLR cells, see Experimental Procedures) for 48 hr and then isolated by FACS based on Pax3-driven GFP expression. Gene expression and chromatin accessibility were concomitantly evaluated in the same cell by single cell nuclear RNA-seq (snRNA-seq) and scATAC-seq multiome. Individual analysis and visual inspection of snRNA-seq and scATAC-seq datasets indicated high concordance of the two approaches in the identification of specific cell clusters (Figure 3A, snRNA and scATAC panels). Next, we integrated snRNA-seq and scATC-seq measurements by weighted-nearest neighbor (WNN) analysis (Figure 3A, WNN panel). WNN is an unsupervised framework that allows to learn the relative utility of simultaneous measurement of multiple modalities (Hao et al., 2021). Clusters obtained by WNN analysis were resolved to generate developmental trajectories (Figure 3B). To compute pseudotemporal trajectories, it is necessary to establish the start (root) of the trajectory. Based on WNN analysis, we placed cells with aPSM gene signatures at the root of the trajectory (Figure 3B, Supplemental Table S3). Pseudotemporal ordering identified clusters which were subsequently integrated with WNN (Figure 3C). Binding of transcription factors (TFs) to *cis*-regulatory DNA sequences controls gene expression programs that define cell state and lineages (Davidson, 2006). Therefore, determining simultaneous expression of TFs and enrichment of their cognate DNA binding motifs (DBMs) should aid in establishing cell identity. Anterior PSM TFs Meis1 and Pbx1 were broadly expressed and enriched in root clusters (Supplemental Figure S3C). When unbiasedly queried, the accessible chromatin of root clusters displayed overrepresentation of Meis/Pbx DBMs (Supplemental Figure S3C). Co-expression of Meis and Pbx and enrichment of their cognate DBMs are consistent with the observation that Meis and Pbx heterodimerize and facilitate chromatin interaction with additional TFs (Knoepfler et al., 1997) (Knoepfler et al., 1999) (Berkes et al., 2004) (Dell’Orso et al., 2016). Sixteen (16) TFs, expressed in clusters pseudotemporally juxtaposed to the root cluster, are involved in regulating pattern specific processes (Supplemental Table S3). Among these TFs, Fli1, which is required for angiogenesis, was also expressed with the vessel-related genes Flt1 and Flt4 in a small cluster (Figure 3A,C vessels cluster). After emerging from root clusters, the trajectory bifurcated to end at two distantly located pseudotemporal clusters (Figure 3B,C). Neurogenic basic-helix-loop-helic (bHLH) Ascl1 and Nhlh1 were expressed and their DBMs overrepresented and footprinted in cluster 8 (Figure 3A,C neurogenesis cluster, and D). Sox2, the bHLH NeuroD4, Pax2, Pax8, and Lhx5 were also expressed in this cluster (Supplemental Figure S3D,E). The overall gene program of cells hosted in this cluster is indicative of neuronal maturation (neuron projection development, synaptic signaling and assembly) (Supplemental Table S3). Lhx5, Pax2, and Pax8 establish a GABAergic inhibitory neurotrasmitter program in the mouse (Pillai et al., 2007).

**Figure 3.**
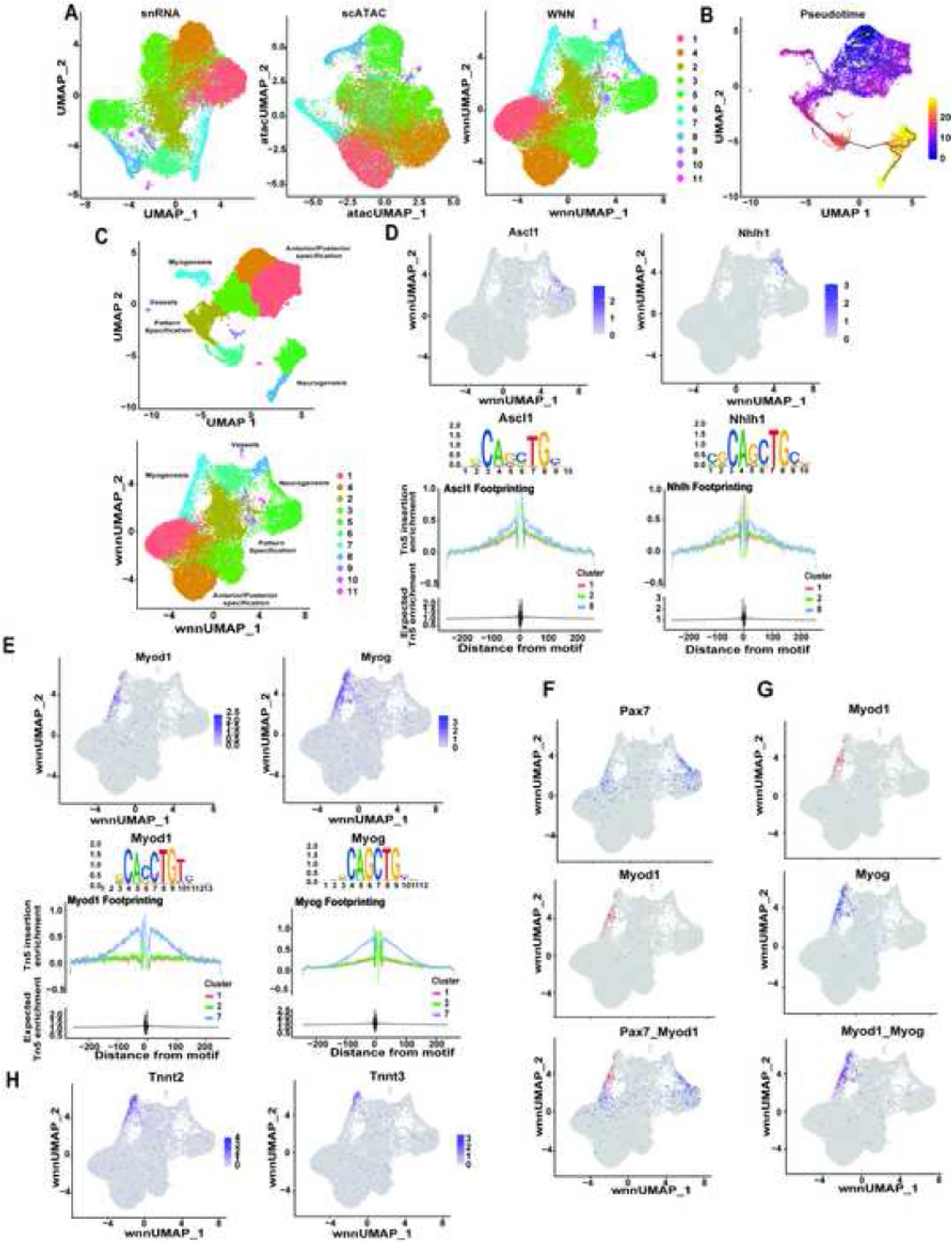
Single Cell Omics of ESC-Derived Cell Lineages. (A) HIFLR cells clustering based on RNA, ATAC, and WNN analysis. (B) Clusters trajectory inference (pseudotime). (C) Clustering derived from trajectory inference (top panel) and clusters identified from trajectory inference projected on WNN (bottom panel). (D) Expression, DNA binding motifs, and footprinting for Ascl1 and Nhlh1 (top to bottom). (E) Expression, DNA binding motifs, and footprinting for Myod1 and Myogenin (top to bottom). (F) Paired-plots expression of Pax7 and Myod1. (G) Paired-plots expression of Myod1 and Myogenin. (H) Tnnt2 and Tnnt3 expression.

The other branch of the trajectory ended at a cluster (Figure 3A,C myogenesis cluster) which hosted cells expressing myogenic bHLH Myod1, Myog, and Myf5 (Figure 3E, Supplemental Figure S3F, and Supplemental Table S3). In this cluster, there was an overrepresentation of DBMs for Myod1, Myog, and Myf5 and footprinting analysis indicated that these DBMs were occupied (Figure 3E, Supplemental Figure S3F). Pax7 directs transcription in neuronal precursors and MuSCs (McKinnell et al., 2008) (Blake and Ziman, 2014) (Lilja et al., 2017) (Magli et al., 2019) (Zhang et al., 2020). In addition to neurogenic cluster 5, Pax7^+^ cells resided in an area transitioning from cluster 1 to cluster 7 where they coexisted with Myod1^+^ cells (Figure 3F). Cells expressing Myod1 or Myogenin were located in the middle section of cluster 7 with Myogenin-positive cells extending towards the tip of the cluster (Figure 3G). The tip of cluster 7 hosted cells expressing the muscle structural troponin Tnnt2 and Tnnt3, and was devoid of Myod1^+^ cells (Figure 3F-H). Thus, cluster 7 may recapitulate *in vivo* skeletal myogenesis where Pax7^+^/Myod1^+^ MuSCs cells progressively extinguish Pax7 while maintaining Myod1 expression (activated MuSCs and proliferating myoblasts, Pax7^-^/Myod1^+^), initiate to differentiate (myocytes, Pax7^-^/Myod1^+^/Myogenin^+^), and terminally differentiate (Myod1^-^/Myogenin^-^/Tnnt3^+^).

### Identification and Characterization of Genomic Regions Regulating Pax7 Expression in Induced ESCs and Myogenic Lineage

Pax7 regulates neuronal gene expression, including spinal cord neurogenesis, and is critical for maintenance and normal function of MuSCs (Mansouri and Gruss, 1998) (Seale et al., 2000) (Oustanina et al., 2004) (von Maltzahn et al., 2013). As chromatin accessibility can predict gene activation, we inspected the Pax7 locus in ATAC-seq datasets generated at different stages of ESCs differentiation. In addition, we wished to compare chromatin accessibility of the Pax7 locus in ESCs and in *bona fide* MuSCs and muscle precursors. For this, we performed ATAC-seq from freshly FACS-isolated MuSCs and from FACS-isolated GFP^+^ cells from E12.5 embryo dissected somites obtained by crossing constitutive Pax7^+^/Cre; Rosa26-YFP mice Compared to naïve and instructed ESCs, aPSM cells displayed increased Pax7 chromatin accessibility, which further augmented in HIFLR cells (Figure 4A). The Pax7 promoter and several regions, including those located at −25Kb, −3.5Kb from the TSS, and in the 7^th^ intron (E7), were also accessible in MuSCs and somites (Figure 4A). The transcriptomes of HIFLR cells and E12.5 Pax7-YFP^+^ somites isolated displayed significant correlation (R^2^>0.8) (Supplemental Figure S4A). scATAC-seq identified a main cluster (cluster 0) of aPSM cells displaying preferential chromatin accessibility at genes related to anterior/posterior pattern specification and somitogenesis (Figure 4B and Supplemental Figure S4B, cluster 0). In HIFLR cells, the main cluster (cluster 1) was enriched for accessibility to genes related to muscle structure development (Figure 4B and Supplemental Figure S4C). Two minor scATAC clusters (clusters 3 and 4) were enriched for accessibility at neurogenic and blood vessel genes in both aPSM and HIFLR cells, respectively (Figure 4B and Supplemental Figure S4B,C). Pax7 was accessible only in the HIFLR myogenic and neurogenic clusters (Figure 4C). Often, when activated, enhancers establish physical contact with the gene promoters that they regulate. We performed *in situ* chromosome conformation capture (Hi-C) (Rao et al., 2014) in aPSM and HIFLR cells and compared them with Hi-C of pluripotent ESCs (Kurup et al., 2020). Hi-C analysis revealed that the Pax7 gene is positioned in a topologically associating domain (TAD) (Supplemental Figure S4D). Despite higher coverage of chromatin interactions, contacts between the Pax7 promoter and E7 element were not detected in pluripotent ESCs. Instead, the E7 region looped to interact with the Pax7 promoter region in both aPSM and HIFLR cells (Figure 4D), indicating that promoter selection by E7 precedes transcriptional activation. To evaluate their role in a muscle environment, we sought to interfere with endogenous Pax7 −25Kb, −3.5Kb, or E7 elements in myogenic C2C12 cells by targeting them with the repressor dCas9-KRAB and specific gRNAs. dCas9-KRAB-mediated perturbation of E7, but not of the −25Kb or −3.5Kb element, reduced expression of endogenous Pax7 to levels comparable to those caused by repressing the Pax7 promoter region (Figure 4E). Next, we tested whether E7 behaves as a functional enhancer by cloning it in a luciferase reporter bearing a minimal promoter and stably integrating the resulting construct in the genome of myogenic C2C12 cells. As control, we inserted a genomic region devoid of ATAC-seq peaks, of comparable length, and located −30Kb from the Pax7 TSS into the luciferase reporter. Compared to either promoter alone or control reporter, Pax7 E7 elicited robust luciferase activity (Figure 4F), indicating that this genomic region functions as an enhancer. Finally, we interrogated the functional relevance of these DNA elements in directing Pax7 activation by deleting the corresponding accessible (ATAC-seq-defined) regions using a CRISPR/Cas9-based approach. Individual or combined biallelic deletion of the −25Kb or −3.5Kb regions in ESCs was without consequence on Pax7 expression (Supplemental Figure S4E,F). In contrast, biallelic deletion of E7 caused a significant reduction (∼70%) of Pax7 expression (Figure 4G). These observations were confirmed in two additional independently isolated ESC clones (Supplemental Figure S4G). Pax7, MyoD, and Myogenin proteins were reduced in E7 deleted cells (Figure 4H). E7 deletion did not interfere with correct Pax7 mRNA splicing (data not shown). We further evaluated the role of E7 by conducting RNA-seq in wild-type and E7 deleted cells. The results of these experiments revealed that activation of the myogenic and neurogenic programs was defective in E7 deleted cells (Figure 4I, Supplemental Figure S4I). Thus, Pax7 E7 controls Pax7 expression and downstream myogenic and neurogenic programs in differentiating ESCs.

**Figure 4.**
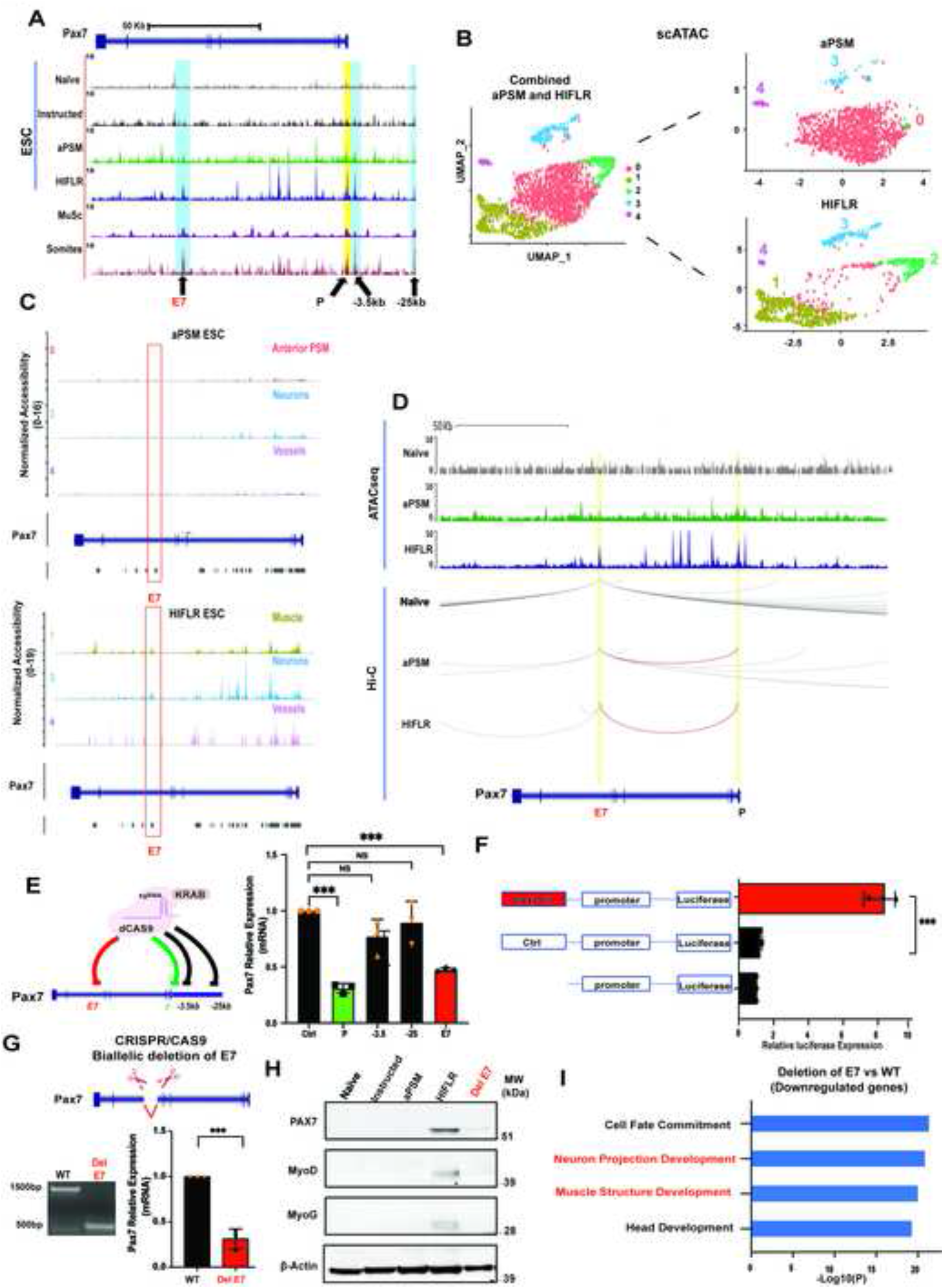
Identification and Characterization of Genomic Regions Regulating Pax7 Expression. (A) Genome browser representation of ATAC-seq tracks at the Pax7 locus in naïve, instructed RDL, HIFLR, MuSCs, and E12.5 YFP^+^ dissected somites from Pax7^+^/Cre;Rosa26-YFP mice. E7 indicates a region located within the Pax7 7^th^ intron; P, promoter; −3.5Kb and −25Kb, regions upstream the Pax7 TSS. (B) scATAC-seq clusters of aPSM and HIFLR cells (left panel, combined clusters; right panel, aPSM and HIFLR individual clusters). (C) scATAC-seq tracks at the Pax7 locus in aPSM (top panel) and HIFLR (bottom panel) clusters. (D) ATAC-seq tracks and Hi-C interactions at the Pax7 locus in pluripotent (ESCs), aPSM, and HIFRL cells. The red line indicates E7-promoter interaction. (E) Scheme representing gRNA-mediated dCas9-KRAB targeting of the indicated Pax7 regions in myogenic C2C12 cells (left panel). Quantitative PCR was employed to measure Pax7 mRNA in myogenic C2C12 cells transfected with dCas9-KRAB and specific or control gRNAs (right panel) Data are represented as mean −/+SD (n=3). Significance is displayed as p<0.001 (***). (F) Luciferase assay in C2C12 cells transfected with the indicated reporter constructs. Data are represented as mean −/+SD (n=3). Significance is displayed as p<0.001 (***). (G) Scheme representing biallelic E7 deletion (top panel). DNA electrophoresis documenting biallelic E7 deletion. Quantitative PCR was employed to measure Pax7 mRNA in control and E7-deleted HIFLR cells (bottom panel). Data are represented as mean −/+SD (n=3). Significance is displayed as p<0.001 (***). (H) Immunoblot of Pax7, MyoD, Myogenin, and beta-actin in in nave, instructed, aPSM, HIFLR, and HIFLR DelE7cells. (I) RNA-seq derived gene ontology terms of downregulated transcripts in E7 deleted HIFLR cells.

## DISCUSSION

This study contributes a bulk well as single-cell resolution resource of both transcriptomes and chromatin accessibility of ESCs induced to differentiate to acquire a PSM-like fate and initial myogenesis and neurogenesis. Exposure to Bmp4 activated gene subsets expressed in ESCs transitioning from naïve to primed pluripotency, a state defined as formative pluripotency. Enhancers and promoters employed different syntaxes to activate or repress ESCs induced genes with chromatin accessibility often preceding gene expression or perduring even when transcription subsided. Single cell approaches permitted the identification of heterogeneous cell types including myogenic and neurogenic precursors. Gene signatures detected in instructed ESCs suggest that neuromesodermal progenitors may participate to the specification of both neurogenic and myogenic precursors (Henrique et al., 2015) (Gouti *et al*., 2017). Neural progenitors and differentiated neurons were observed during the initial phases of directed myogenic differentiation of human pluripotent stem cells (Xi *et al*., 2020). Underlying species difference, neurogenic and myogenic cells emerged 3 and 4 weeks after directed differentiation of human pluripotent stem cells (hiPSCs), respectively (Xi *et al*., 2020) whereas data reported here indicate that mouse pluripotent stem cells give rise to neurogenic and myogenic cells 8 days after initial ESCs induction. In two other studies, hiPSCs activated the myogenic program after 10-15 days, though with different PAX7 activation kinetics (Wu et al., 2018) (Al Tanoury *et al*., 2020). The differences in neurogenic and myogenic induction and myogenic activation kinetics are likely due to different protocols employed in different studies. Given the relevant role played by Pax7 in neurogenesis and myogenesis, we have leveraged our datasets to identify Pax7 regulatory regions directing expression in both cell lineages. The enhancer region we have identified is distinct from those directing Pax7 expression in mouse cranial neural crest, facial mesenchyme, mesencephalon, pontine reticular nucleus (Lang et al., 2003) and from the enhancer that directs Pax7 expression in chick embryos neural crest (Vadasz et al., 2013). We anticipate that integration of the datasets reported here to characterize Pax7 regulatory regions may be employed to query the dynamics of gene regulatory networks occurring in ESCs induced to acquire a PSM-like fate, initial myogenic or neurogenic differentiation.

### Limitations of the study

Mouse ESCs represent a culture model and do not reflect the complex series of events occurring during development. Transcriptomes and epigenomes of ESCs only partially recapitulate those observed in the embryo. Moreover, our study is limited to four ESC time-points (naïve ESCs, Bmp4-instructed ESCs, aPSM, and HIFLR cells). A more detailed time-course evaluation of ESCs induced to acquire a presomitic mesoderm (PSM) fate and longer exposure of PSM cells to differentiating factors would add further granularity to our datasets. An additional limitation of this study relates to the composition, doses, and timing of cells exposure to defined factors and small molecules which determine the outcome of the observations.

## Experimental Model and Subject Details

### Cell lines

All cells were cultured at 37°C with 5% CO_2_. HEK293T and C2C12 cells (ATCC) were grown in 1× DMEM supplemented with 10% and 20% of qualified fetal bovine serum (FBS) (GIBCO), respectively.

Mouse Embryonic Stem Cells (ESCs) expressing Pax3-GFP are described in (Chal et al., 2016) ESCs were cultured as follow:

*Maintenance.* ESCs were plated on 0.1% gelatin (Millipore) and cultured in DMEM (Gibco) supplemented with 15% inactivated fetal bovine serum (Hyclone), 1% penicillin-streptomycin, 2 mM L-glutamine, 0.1 mM non-essential amino acids, 0.1% β-mercaptoethanol, 1,500 U/ml LIF and 2i inhibitors (Stemgent).

*Serum-free differentiation of mouse ESCs.* ESCs differentiation was induced as described in Chal *et al*., 2015 with minor modifications. Briefly, ESCs were trypsinized and plated on gelatin-coated plates in serum-free N2B27 medium (N2B27,1% Knock-out Serum Replacement (KSR, Gibco), 0.1% bovine serum albumin (Gibco) and BMP4 (Peprotech) at 10 ng/ml. After 2 days, cells were shifted to RDL medium, (DMEM, 1%FBS, 14% KSR,10 ng/ml Rspo3 (Peprotech, R&D Biosystems), 0.5% DMSO (Sigma), and 0.1 μM LDN193189 (Stemgent), and cultured for 4 additional days. Subsequently, medium was changed to DMEM, 1%FBS, 14% KSR, 0.1% BSA supplemented with 10 ng/ml Rspo3, 10 ng/ml HGF, 2 ng/ml IGF-1, 20 ng/ml FGF-2 (Peprotech, R&D Biosystems) and 0.1 μM LDN193189 (HIFLR medium).

### Mice and animal care

Mice were housed in a pathogen-free facility, all the experiments were performed according to the National Institutes of Health (NIH) Animal Care and Use regulations. Pax7^+^/Cre; Rosa26-YFP embryos were generated by timed mating between Pax7^+^/Cre (Keller et al., 2004) and Rosa26-YFP (Rosa26-YFPY/Y) mice (Srinivas et al., 2001).

### Bacterial strains

DH5α and TOP10 cells were obtained from New England BioLabs (NEB) and Thermo Fisher Scientific Inc. Both DH5α and TOP10 bacterial strains were grown in LB medium at 37°C and used to propagate plasmids.

## Methods Details

### Single guide RNA (sgRNA) design and Cas9 vector assembly

sgRNA for CRISPR/Cas9 assay were designed using *CHOPCHOP* web tool (https://chopchop.cbu.uib.no/). sgRNAs were tested by T7 Endonuclease assay (https://www.neb.com/products/m0302-t7-endonuclease) to evaluate editing efficiency, and selected sgRNAs were cloned into the Cas9 vector pSpCas9 (BB)-2A-GFP (pX458; Addgene plasmid#48138) or into the dCAS9-KRAB repressor GFP-plasmid (addgene#71236) (Primers are reported in Supplemental Table S4).

#### Plasmids Construction

The genomic region corresponding to the E7 enhancer (chr4:139771035-139771985) and a control region (chr4:139,863,500-139,864,500.) were amplified and cloned into the 5’ XhoI and 3’ EcorV sites in pGL4.26[luc2/minP/Hygro] vector (Promega, #E8441). Primers are reported in Supplemental Table S4).

### Transfections and Genome Editing

Plasmid transient transfections were performed using Lipofectamine 2000 (Invitrogen). For genome editing, ESCs cells were transfected in suspension with 1.5µg of pSpCAS9-GFP with different sgRNAs and plated on 6 well gelatin-coated plates. 48 hours after transfection cells were seeded in 100mm culture dishes at a density of 1.5 and 10 cells/ml; 2 weeks later, clones were collected and screened for genomic editing. For luciferase reporter assay, 2X10^5^ C2C12 cells were transfected with ∼1-3 µg of Pax7 luciferase vectors and plated in 6 well plates. 72 h post-transfection medium was changed and cells were selected with 300μg/ml Hygromycin (Sigma, 31282-04-9) for 5 days.

### Lentiviral Production

Lentiviral particles were generated using HEK293T cells (ATCC). Cells were seeded at 50-60% confluency in 100 mm cell culture dish. The following day, cells were transfected with 5μg of the vector of interest (dCAS9-KRAB with specific sgRNAs), 1μg of pMD2.G and 4μg of psPAX2 (Addgene) using 30μl of Lipofectamine 2000). 24 hr post-transfection the media was changed. The supernatant was collected 48hr post-transfection, filtered with a 0.45-mm PVDF filter (Millipore), and concentrated using the lenti-x concentrator (Takara, 631232) according to the manufactorer’s instructions. Concentrated viruses were stored at −80C. C2C12 cells were transduced in suspension and plated in DMEM medium supplemented with 6 μg polybrene. Cells were collected five days post-transduction and GFP positive cells were sorted for RNA extraction.

### Luciferase Reporter Assay

Luciferase activity was detected by adding luciferin (Promega, E1500) to cell extracts of transfected C2C12 cells as a substrate at a final concentration of 0.15 mg/mL according to the manufacturer’s instructions. Fluorescence was detected and quantified on a luminometer (Synergy™ H1 microplate reader, BioTek). Luciferase activity was normalized by total protein concentration.

#### ESCs FACS Sorting

ESCs were trypsinized and FACS-isolated gating on positive GFP and DAPI staining, and negative PI staining using FACSAria Fusion (BD Biosciences).

#### Somites Isolation and FACS Sorting

Somites were isolated from E12.5 Pax7^+^/Cre; Rosa26-YFP embryos. Cells were isolated from regions corresponding to posterior to forelimb area or mid-trunk area. 20 to 25 pairs of somites were obtained from each embryo. The spinal cord and lateral paraxial mesoderm were separated by sharp forceps to avoid sorting of YFP^+^ dorsal spinal cord cells. The separated lateral tissue was immediately placed into 500ul of 2.5mg/ml Papain/1xPBS (Sigma-Aldrich), minced with scissors and incubated at 33°C for 20 minutes for disassociation. 500ul of sorting medium (15% FBS in 1x PBS) was added to the digested tissue to stop the reaction. The obtained tissue was diluted in 10ml of sorting medium (15% FBS in 1x PBS), then filtered with a 0.45mm filter (Millipore). Cells were centrifuged at 1300 rpkm for 5 min and resuspended in 500ul of Sorting medium. Final FACS-sorting was conducted on a BD FACSAria IIIu machine by gating the YFP channel. About 100,000-200,000 YFP^+^ somitic cells were collected from one sorted embryo.

#### Muscle Dissection and MuSC FACS-Isolation

Hindlimb muscles from 3-month-old C56BL/6J wild-type mice were dissociated into single cells by enzymatic digestion and live cells were isolated by FACS. Skeletal MuSCs were sorted following described methods (Liu et al., 2015). Briefly, hindlimb muscles from 3-month-old adult wild-type mice were minced and digested with collagenase for 1 h and MuSCs were released from muscle fibers by further digesting the muscle slurry with collagenase/dispase for an additional 30 min. After filtering out the debris, cells were incubated with the following primary antibodies: biotin anti-mouse CD106 (anti-VCAM1, BioLegend 105704; 1:75), PE/Cy7 Streptavidin (BioLegend 405206; 1:75), Pacific Blue anti-mouse Ly-6A/E (anti-Sca1, BioLegend 108120; 1:75), APC anti-mouse CD31 (BioLegend 102510; 1:75) and APC anti-mouse CD45 (BioLegend 103112; 1:75). Satellite cells were sorted by gating VCAM1-positive, Pacific Blue-labeled Sca1-negative, and APC-labeled CD31/CD45-negative cells. SYTOX Green (ThermoFisher Scientific S7020; 1:30,000) was used as a counterstain

### Antibodies

Western blot experiments were conducted using the following antibodies: anti-Pax7 (DSHB), anti-Myod1 (Santa Cruz. 71629), anti-MyoG (Santa Cruz. 12732), anti-b-Actin (Santa Cruz. 130656), anti-Otx2 (cell signaling. mAb #11943), and anti-Nanog (Santa Cruz, sc-374001). Goat anti-Rabbit IgG-HRP or goat anti-mouse IgG-HRP (Azure biosystem, AC2114 and 2115 respectively) were used as secondary antibodies for immunoblotting. Antibodies anti-H3K27Ac (ab4729, Abcam) and anti-H3K4me1(ab8895, Abcam) were employed in ChIP-seq experiments.

### Quantitive and Semi-quantitive PCR

Total RNA was extracted using TRIzol Reagent (Invitrogen) according to the manufacturer’s protocol. cDNAs were synthesized with the qScript cDNA kit (Quanta) containing random primers. Reverse transcribed cDNA was diluted 10 times and SYBR green real-time PCR was performed on the Applied Biosystems StepOne Plus Real-Time PCR system. Quantifed mRNA levels were normalized to 18S and relative expressions were calculated according to ΔΔCt. A full list of primers is reported in Supplemental Table S4.

To screen for positive CRISPR/CAS9 clones carrying the expected deletions, genomic DNA was extracted using a Blood & Tissue Extraction kit (Qiagen), and PCR was performed on Applied Biosystems PCR system. Primers are reported in Supplemental Table S4.

### Western Blotting

Cells were lysed in lysis buffer [100 mM Tris-HCl pH 7.5 and 5% sodium dodecyl sulfate (SDS)]. Proteins (20-30 μg) were incubated at 95 °C for 5 minutes, resolved by SDS-PAGE, and transferred to nitrocellulose membranes. Subsequently, membranes were incubated with blocking solution (0.1% Tween 20, 5% non-fat milk in 1XPBS) for 1 h at room temperature, and primary antibodies were added overnight at 4 °C. Membranes were then washed with 1XPBS/0.05% Tween 20 and incubated with secondary antibody (Azura HRP Ab, CatN X) for 1 h at room temperature. Bioanalytical Imaging System-600 (Azura Biosystem, Inc) was used for protein visualization.

### ChIP-Seq

4 × 10^6^ ESCs were used for chromatin immunoprecipitation. Cells were crosslinked in 1% formaldehyde and processed according to published protocols (Métivier et al., 2003, Mousavi et al., 2012). Briefly, cells were lysed in RIPA buffer (1× PBS, 1% NP-40, 0.5% sodium deoxycholate, 0.1% SDS) and centrifuged at 2,000 rpm for 5 min. The chromatin fraction was shared by sonication (four times, each lasting 30). The resulting sheared chromatin samples were immunoprecipitated overnight, and washed in buffer I (20 mM Tris-HCl [pH 8.0], 150 mM NaCl, 2 mM EDTA, 0.1% SDS, 1% Triton X-100), buffer II (20 mM Tris-HCl [pH 8.0], 500 mM NaCl, 2 mM EDTA, 0.1% SDS, 1% Triton X-100), buffer III (10 mM Tris-HCl [pH 8.0] 250 mM LiCl, 1% NP-40; 1% sodium deoxycholate, 1 mM EDTA), and Tris-EDTA (pH 8.0). All washes were performed at 4°C for 5 min. Finally, crosslinking was reversed in elution buffer (100 mM NaHCO3, 1% SDS) at 65°C overnight. For ChIP-seq, 10 ng immuno-precipitated DNA fragments were used to prepare ChIP-seq libraries with the NEBNext Ultra II DNA library prep kit for Illumina (New England Biolabs, (#E7645) following the manufacturer’s protocol. The libraries were sequenced on a NextSeq550 or NovaSeq6000 Illumina instrument.

### ATAC-Seq

ATAC-seq was performed according to a published protocol (Buenrostro et al., 2013) with minor modifications. Briefly, 5×10^4^ cells were washed with 50ul of 1xPBS and lysed in 50ul of Lysis Buffer (10mM Tris-HCl, pH7.4, 10mM NaCl, 3mM MgCl2, 0.1% of IGEPAL CA-630). To tag and fragment accessible chromatin, nuclei were centrifuged at 500x g for 10min and resuspended in 40ul of transposition reaction mix with 2ul Tn5 transposase (Illumina Cat# FC-121-1030). The reaction was incubated at 37°C with shaking at 300rpm for 30min. DNA fragments were then purified and amplified by PCR (12–15 cycles based on the amplification curve). Purified libraries were sequenced on NextSeq550 or NovaSeq6000 Illumina instrument.

### RNA-Seq

For transcriptome analysis (RNA-seq), poly (A)^+^ mRNA libraries were generated in triplicate using NEBNext Ultra II RNA library preparation kit for Illumina (NEB #E7490) according to the manufactory instructions.

### Single Cell RNA-Seq

ESCs were collected, washed once with 2×PBS, and re-suspended in PBS with 0.04% bovine serum albumin. Cellular suspensions were loaded on a Chromium Instrument (10x Genomics) to generate single-cell GEMs. Single-cell RNA-seq libraries were prepared using a Chromium Single Cell 3’ Library & Gel Bead Kit v3.1 (P/N 1000121, 10x Genomics). GEM-RT was performed in a C1000 Touch Thermal cycler with 96-Deep Well Reaction Module (Bio-Rad; P/N 1851197): 53°C for 45 min, 85°C for 5 min; held at 4°C. Following retrotranscription, GEMs were broken and the single-strand cDNA was purified with DynaBeads MyOne Silane Beads. cDNA was amplified using the C1000 Touch Thermal cycler with 96-Deep Well Reaction Module: 98°C for 3 min; cycled 12 times: 98°C for 15 s, 63°C for 20 s, and 72°C for 1 min; 72°C for 1 min; held at 4°C. Amplified cDNA product was purified with the SPRIselect Reagent Kit (0.6× SPRI). Indexed sequencing libraries were constructed using the reagents in the Chromium Single Cell 3’ Library & Gel Bead Kit v2, following these steps: (1) end repair and A-tailing; (2) adaptor ligation; (3) post-fragmentation, end repair, and A-tailing double size selection cleanup with SPRI-select beads; (4) sample index PCR and cleanup. The barcode sequencing libraries were diluted at 3 nM and sequenced on an Illumina NovaSeq6000 using the following read length: 28bp for Read1, 8 bp for I7 Index, and 91 bp for Read2.

### Single Cell ATAC-Seq

Cells were washed twice with 1xPBS with 0.04%BSA (Sigma) and single-cell ATAC was performed using Chromium Next GEM Single Cell ATAC Library & Gel Bead Kit v1.1 (10x genomics, 1000175). Briefly, 1X106 cells were lysed using chilled lysis buffer (10mM Tris HCl, 10mM NaCl, 3mM MgCl2, 0.1% Tween 20, 0.1% Nonidet P40 substitute, 0.01% Digitonin and 10% BSA). Following lysis, nuclei were washed with wash buffer (10mM Tris HCl, 10mM NaCl, 3mM MgCl2, 0.1% Tween 20 and 1% BSA) and re-suspended in Nuclei buffer (10x Genomics, PN-2000153). Subsequently, nuclei suspensions were incubated with transposition mix for 1 hour in 37C in a C1000 Touch Thermal cycler. Nuclei were diluted in Diluted Nuclei Buffer according to 10X Genomics recommendations (10X Genomics) and loaded on a Chromium Instrument (10x Genomics) to generate GEMs. Samples were incubated in a C1000 Touch Thermal cycler with 96-Deep Well Reaction Module (Bio-Rad; P/N 1851197) programmed as 72°C for 5 min; 98°C for 30 s, cycled 12 times: 98°C for 10 s, and 59°C for 30s; 72°C for 1 min; held at 4°C. Amplified products were purified with Dynabeads MyOne Silane beads followed by SPRIselect Clean up. Indexed sequencing libraries were constructed using the reagents in the Chromium Single Cell ATAC Reagent Kit v1.1 to add the P7 and P5 sequences used in Illumina bridge amplification, and a sample index. Final libraries were diluted to 3 nM and sequenced on an Illumina NovaSeq6000 using the following read length: 50 bp for Read1, 8 bp for I7 Index, 15 bp for i5 Index, and 50 bp for Read2.

### Chromosome Conformation Capture-Hi-C

Hi-C experiments were performed using the Arima-HiC kit (A510008GFP). Briefly, 2×10^6^ cells were crosslinked with 2% formaldehyde (37%, Sigma) for 10 min followed by incubation with 200mM glycine (sigma) to stop cross-linking (sigma). Cells were washed three times with 1xPBS and resuspended in 20ul elution buffer (Arima). Cells were then lysed and chromatin was extracted, digested, and biotinylated following Arima-HiC kit instructions. Biotin-labeled DNA was quantified and fragmented to 400 bp length using Covaris (ME220) About 300ng of DNA was used for biotin enrichment. Final DNA libraries were constructed using the Swift Biosciences kit for Library Preparation (Accel-NGSO 2S Plus DNA Doc A160140 v00).

### Single Cell Multiome

Single-cell ATACseq and gene expression omics were performed using Chromium Next GEM Single Cell Multiome ATAC+ Gene Expression (10x Genomics kit, CG000338) according to the manufacturer’s instruction. Briefly, cells were washed with 1xPBS and resuspended in PBS-0.04%BSA. Nuclei were isolated by incubating the cells in chilled Lysis Buffer (10mM Tris HCl, 10mM NaCl, 3mM MgCl2, 0.1% Tween 20, 0.1% Nonidet P40 substitute, 0.01% Digitonin and 10% BSA) for 3 minutes. After washing, nuclei suspensions were incubated in a Transposition Mix that includes a Transposase and loaded into the Chromium instrument (10X Genomics) for GEMs generation. Samples were then incubated in a C1000 Touch Thermal cycler with 96-Deep Well Reaction Module (Bio-Rad; P/N 1851197) programmed as 37°C for 45 min; 25°C for 30 min, 4°C holds. Amplified products were purified with Dynabeads MyOne Silane beads followed by SPRIselect clean up. Eluted samples containing barcoded transposed DNA and barcoded cDNA were pre-amplified via PCR and cleaned up using SPRIselect. Eluted pre-amplified reactions were used to generate ATACseq and gene expression (GEX) libraries according to the 10X Genomics protocol. 1μl of each library was run on High sensitivity D1000 TapeStation to determine the correct average fragment size. Libraries were diluted at 3 nM and sequenced on an Illumina NovaSeq6000 using the following read length: 50bp for Read1, 8 bp for I7 Index, 24bp for i5 index, and 49bp for Read2.

### Gene Ontology Analysis

Gene Ontology (GO) analysis of genes from different experiments was performed with Metascape, gene annotation, and analysis resource. (https://metascape.org/gp/index.html#/main/step1)

### Bulk RNA-Seq Analysis

RNA-Seq data were generated using Illumina NovaSeq6000 system. Raw sequencing data were processed with bcl2fastq/2.20.0 to generate FastQ files. Adapter sequences were removed using trimgalore/0.6.6(https://github.com/FelixKrueger/TrimGalore). Single-end reads of 50 bases were mapped to the mouse transcriptome and genome mm10 using TopHat 2.1.1 (Trapnell et al., 2009). Gene expression values (RPKM: Reads Per Kilobase exon per Million mapped reads) were calculated using Partek Genomics Suite 7.18, which was also used for the PCA and ANOVA analyses.

### ChIP-Seq and ATAC-Seq Analysis

Sequencing data were generated with an Illumina NovaSeq6000 system. FastQ files were generated with bcl2fastq/2.20.0. Adapter sequences were removed using trimgalore/0.6.6. Reads of 50 bases were aligned to the mouse genome build mm10 with Bowtie/1.1.1 (Langmead et al., 2009), allowing two mismatches. Uniquely mapped and non-redundant reads were used for peak calling using MACS 1.4.2 (Zhang et al., 2008) with a p value cutoff of 1.0E-05. Histone model was applied for CHIP-seq samples while transcription factor model for ATAC-seq samples. Only regions called in both replicates were used in downstream analysis. Bigwig files were generated with BedGraphToBigWig (Kent et al., 2010)and Bedtools/2.29.2 (Quinlan and Hall, 2010). Peak intensities were normalized as tags per 10 million reads (RP10M) in the original library. Peaks were assigned to the closest TSS with HOMER.

### Single Cell RNA-Seq Analysis

Demultiplexing and reads alignement were performed with CellRanger ver. 3.1.0 (10x Genomics) with default parameters. For all analyses, we employed standard pre-processing for all single-cell RNA-seq datasets. Filtering, variable gene selection, and dimensionality reduction were performed using the Seurat ver.4.1.0 (Butler et al., 2018).

#### Filtering

1. Cells were filtered out based on the following criteria:

- To eliminate low quality cells and debris, cells with less than 200 detected genes and cells with more than 20% of UMIs mapped to mitochondrial genes were excluded.
- To remove potential cell doublets and aggregates only cells showing an nFeature_RNA between 1600 and 8000 were selected.

#### Normalization

1. For each cell, UMI counts per million were log-normalized using the natural logarithm.
2. In each dataset, we aimed to identify a subset of features (e.g., genes) exhibiting high variability across cell to prioritize downstream analysis. Variable genes were selected applying thresholds calculated using binned values from log average expression and dispersion for each gene.

#### Clustering

1. The expression level of highly variable genes was scaled along each gene and cell-cell variation was regressed out by number of detected molecules, and mitochondrial gene expression.
2. Data were projected onto a low-dimensional subspace of PCA (principal component analysis) using dimensional reduction. The number of PCA were decided through assessment of statistical plots.
3. Cells were clustered using a graph-based clustering approach optimized by the Louvain algorithm with resolution parameters and visualized using two-dimensional UMAP (Uniform Manifold Approximation and Projection).
4. To define cluster identity, expression and distribution of known markers were evaluated.

### Integration of scRNA-Seq Datasets

We integrated scRNA-seq datasets of naïve and instructed ESCs using Harmony (v1.0) (Korsunsky et al., 2019) for batch correction. Harmony was applied on the first 30 PCA components (RunHarmony function setting assay.use to RNA, max.iter.harmony to 10, max.iter.cluster to 20, sigma to 0.1). After dimensional reduction, cells were clustered by the Louvain algorithm (resolution at 0.25,1.0 and 2.0), and gene expression distributions were visualized using two-dimensional UMAP Harmony.

### Pseudotemporal Analyses

Pseudotime was calculated by the Monocle3 R package (Cao et al., 2019) following conversion of gene expression data and cell metadata including cell type labels from the Seurat object to Monocle object. After passing parameters in Seurat to the ‘‘residual_ model_formula_str’’ argument in the ‘‘preprocess_cds’’ function, significant PCs were selected to further reduce the data dimensionality using uniform manifold approximation (UMAP). The beginning of pseudotime was selected in the UMAP dimension and distribution of the pseudotime was calculated by “order_cells” function. Slingshot(ver 2.2.0) (Street et al., 2018) and tradeSeq(ver 1.8.0) (Van den Berge et al., 2020) were used to infer the pseudotemporal ordering along the trajectory of the cellular embedding of the single cell multiome dataset.

### Single Cell ATAC-Seq Analysis

Demultiplexing and reads alignment were performed using Cellranger-atac ver. 1.1.0 with default parameters). For all analyses, we employed standard pre-processing for all single-cell ATAC-seq datasets. Filtering, variable gene selection, and dimensionality reduction were performed using the Signac ver.1.5.0 (Stuart et al., 2019).

#### Filtering

ESCs with fewer than 3 detected genes were excluded. Additional standard QC steps (duplicates removal, evaluation of number of fragments per cell and number of fragments per peak, fraction of reads mapping to blacklist regions, nucleosome pattern signal, and transcriptional start site (TSS) enrichment) were also applied.

#### Clustering

1. LSI dimensionality reduction was applied to all samples (RunTFIDF function with method equal to 2, FindTopFeatures function setting min.cutoff to q0, and RunSVD function using the peaks as assay) and data were projected onto a low-dimensional subspace of PCA (principle component analysis) using dimensional reduction.
2. Cells were clustered using a graph-based clustering approach optimized by the Louvain algorithm with resolution parameters, and visualized using two-dimensional UMAP (Uniform Manifold Approximation and Projection).
3. Gene expression in each cluster was predicted emploiyng GeneActivity function, obtained gene activity was generated in the Seurat object (normalization method with log-normalized and scale factor with the median of the sample).

### Downstream Analysis of scATAC-Seq Data

Downstream analysis was performed using R 4.1.0 applying Signac ver.1.5.0 (Stuart et al., 2021). After correction of GC bias using RegionStats function in Sinac, peaks correlated with the expression of nearby genes were identified with LinkPeaks function. Next, the tracks of ATAC-seq peaks for each cluster were visualized using CoveragePlot function (−0.5Kb/+14Kb from the TSS). Transcriptional factor (TF) activities on the ATAC-seq data were calculated using the Signac implementation of TFBSTools (Tan and Lenhard, 2016) with JASPAR2020 vertebrates TF binding models database (746 TFs) (Fornes, Oriol et al., 2019). The footprinting of each TF near the TSS regions was visualized using Footprint and PlotFootprint functions.

### Integration of scATAC-Seq Data

scATAC-seq datasets were integrated as described (Stuart *et al*., 2019). Common anchors in the scATAC-seqs datasets were identified by Seurat function FindTransferAnchors with Latent Semantic Indexing (LSI) reduction and the first 40 components were calculated. These components were then used to generate a Uniform Manifold Approximation and Projection (UMAP) dimensionality reduction. Nearest-Neighbor graph was generated after excluding the first component following the standard guidelines from the Satija lab (https://satijalab.org/signac/). Cells were then clustered with the Seurat’s Louvain algorithm.

### Integration of scRNA-seq and scATAC-seq Data

scATAC-seq and scRNA-seq data were integrated as described (Stuart *et al*., 2019). Using scRNA-seq and scATAC-seq data, cluster labels in the scATAC-seq dataset were predicted by the Seurat function FindTransferAnchors on the Canonical Correlation Analysis (CCA) space and were trasnferred to the scATAC-seq dataset. This operation used the variable features of the scRNA-seq analysis on the RNA assay as the reference data, and the gene activity matrix of the scATAC-seq analysis as the query data. After transferring the labels through TransferData function, they were merged into the two Seurat objects, and co-visualized with clusters labeled by the scRNAseq cluster labels. Finally, dot-plots visualizing averaged gene expressions and chromatin accessibility were generated by calculating z-scores.

### Single Cell Multiome Analysis

The multi-omic dataset was realigned to mm10 using cellranger-arc version 2.0.0 (10x Genomics). The resulting RNA count matrix was filtered for cells with between 1,200 and 10,000 reads while ATAC peak matrix was filtered with between 100 and 10,000 reads. Next, the WNN graph was constructed to learn the relative utility of each data modality in each cell(https://satijalab.org/seurat/articles/weighted_nearest_neighbor_analysis.html: WNN analysis of 10x Multiome, RNA + ATAC). The R packages Seurat ver.4.1.0 and Signac ver.1.5.0 were used for data scaling, transformation, clustering, dimensionality reduction, differential expression analysis, and most visualizations.

### Hi-C data analysis

Hi-C datasets were processed using Juicer 1.6 (Durand et al., 2016) Raw reads were aligned to the mm10 reference genome using BWA (Li and Durbin, 2010). Reads that aligned to more than two places in the genome were discarded. The remaining aligned reads were filtered based on mapping quality score (MAPQ <30). Contact matrices were generated at different base pair resolutions ranging from 1MB to 5kb. Downstream analysis was performed with HOMER (Heinz et al., 2010). The Hi-C summary output from Juicer was used to generate a paired end tags using the HOMER makeTagDirectory script. Extraction of significant interactions was done using HOMER at 10kb resolution with a 25kb window, by running the HOMER AnalyzeHiC script. ATAC-Seq peak regions for each sample were used to assess overlapping of significant interactions with regions of open chromatin. HiCExplorer 3.6 (Wolff et al., 2020) was employed to visualize significant interactions along with ATAC-seq tracks for each sample at defined regions of interest. The code executed in the described pipeline is available in Key Resources Table of STAR Methods.

### Statistical analysis

Data from quantitive PCR are expressed as mean ± standard error from three different experiments (n=3). Significant differences were analyzed by the two-tailed, unpaired, Student’s t test and values were considered significant at p<0.05.

## SUPPLEMENTAL INFORMATION

**Supplemental Table S1.** Lists of transcripts up- or down-regulated in instructed ESCs.

**Supplemental Table S2.** Lists of *de novo*, accessible, and unmarked enhancer regions in naïve ESCs. Lists of transcripts enriched in naïve and instructed ESCs scRNA-seq clusters. Transcripts enriched in instructed ESCs and E8.5/E9.5 neuromesodermal progenitors.

**Supplemental Table S3.** Summary of scRNA-seq clusters, differential gene transcripts, total cell number, and cell percentage/cluster. Lists of transcripts enriched in HIFLR scRNA-seq clusters

**Supplemental Table S4.** Oligonucleotide sequences of sgRNA, quantitative and semi-quantitative PCR oligos.

## ACKNOWLEDGMENTS

We thank the NIAMS Genomic Technology Section, Biodata Mining and Discovery Section, Flow Cytometry Section, and Light Imaging Section. This study utilized the high-performance computational capabilities of the Helix Systems at the NIH, Bethesda (https://helix.nih.gov/). This work was supported in part by the Intramural Research Program of the NIAMS at the NIH (grants AR041126 and AR041164 to V.S.).

## AUTHOR CONTRIBUTIONS

M.K. performed most of the experiments. J.P. conceived the study, established initial ESC culture conditions and performed and analyzed ATAC-seq and bulk RNA-seq experiments. KDK and KJ analyzed RNA-seq, ATAC-seq, ChIP-seq, scRNA-seq, scATAC-seq and multiomics data. X. F. isolated somites. N.A-L. analyzed Hi-C data. V.C. helped with libraries preparation. P.G. and A.H. provided advice on CRISPR/Cas9 experiments. J. S. assisted with FACS. S.D.O. provided advice for the genomic studies, analyzed data, and supervised the project. V.S. conceived and designed the study, analyzed data, supervised the project, and wrote the manuscript with input from the authors.

## DECLARATION OF INTERESTS

The authors declare no competing interests.

## Supplemental Figure Legends

**Supplemental Figure S1 (related to Figure 1).**

(A) Scheme of ESCs Pax3-GFP cultured in N2B27 medium supplemented with Bmp4 and 1%KSR without LIF and 2i for 48 hr (Instructed). (B) Selected gene ontology terms of upregulated and downregulated transcripts in instructed ESCs. (C) Averaged normalized tag intensities of H3K27ac at naïve and instructed regulatory regions. (D) Genome browser representation of ATAC-seq and ChIP-seq tracks for Pou3f1, Wnt8, Pbx1, Pitx2, Esrrb, Nanog, and Tbx3 loci. (E) Expression of autophagy Ulk1 and Gabarapl2 transcripts in scRNA-seq naïve and instructed ESCs clusters. (F) Expression of Otx2 and Sall2. (G) scATAC-seq of naïve and instructed ESCs clusters. Gata4, Gata6, Hnf1b, and Foxa2 TSS were accessible and footprinted in the indicated instructed cluster. (H) Expression of Gata6 and Foxa2 transcripts (top panel) and overall scATAC-seq Gata6 and Foxa2 footprinting.

**Supplemental Figure S2 (related to Figure 2).**

(A) Scheme indicating differentiating medium conditions to instruct ESCs to acquire an anterior presomitic mesoderm (aPSM) fate. (B) Fgf8 expression in aPSM cells. (C) Expression of vessel endothelial transcripts Flt1, Flt4, and Kdr in aPSM cells. (D) Expression of neuronal transcripts Pax6, Sox2, and Pou3f2 in aPSM cells.

**Supplemental Figure S3 (related to Figure 3)**

(A) Scheme indicating medium conditions to induce aPSM cells to acquire an initial myogenic cell fate. (B) Expression and DNA binding motifs for Meis1 and Pbx1. (C) Fli1, Flt1, and Flt4 expression. (D) Expression and DNA binding motif for Sox2, and Pax2, NueroD4, Pax8, and Lhx5 expression. (E) Expression, DNA binding motif, and footprinting for Myf5.

**Supplemental Figure S4 (related to Figure 4)**

(A) Gene expression correlation plot of somites and HIFLR cells RNA-seq. (B) Selected gene ontology of scATAC-seq clusters for aPSM and (C) for HIFLR cells. (D) Heatmap of contact matrices from Hi-C data indicating that Pax7 locus resides in a topologically associating domain. (E) Gel electrophoresis of genomic DNA documenting biallelic deletion of the −3.5Kb Pax7 region. Quantitative PCR was employed to measure Pax7 mRNA expression in control and two independent biallelic deleted ESC clones (n=3, NS, not significant). (F) Gel electrophoresis of genomic DNA documenting biallelic deletion of the −3.5Kb and −25Kb Pax7 regions. Pax7 mRNA expression in control and two independent biallelic deleted −3.5Kb/−25Kb ESC clones (n=3, NS, not significant). (G) Gel electrophoresis of genomic DNA documenting monoallelic (one clone) and biallelic deletion of the E7 Pax7 region in two independent clones. Quantitative PCR was employed to measure Pax7 mRNA in control and two independent biallelic E7 deleted ESC clones. Data are represented as mean −/+SD (n=3). Significance is displayed as p<0.001 (***). (H) DNA sequencing chromatogram of a E7 deleted ESC clone. (I) RNA-seq tracks of Pax7, MyoD, Myogenin, Neurogenin, Ascl1, and NeuroD4 in control (WT) and E7 deleted ESCs.

## Notes

### Competing Interest Statement

The authors have declared no competing interest.

