## Supplemental Figures for "Transcriptomics, Regulatory Syntax, and Enhancer Identification in Heterogenous Populations of Mesoderm-Induced ESCs at Single-Cell Resolution"

Supplemental Figure 1

Click here to access/download;Supplemental Figure;Supplemental Fig S1.tif

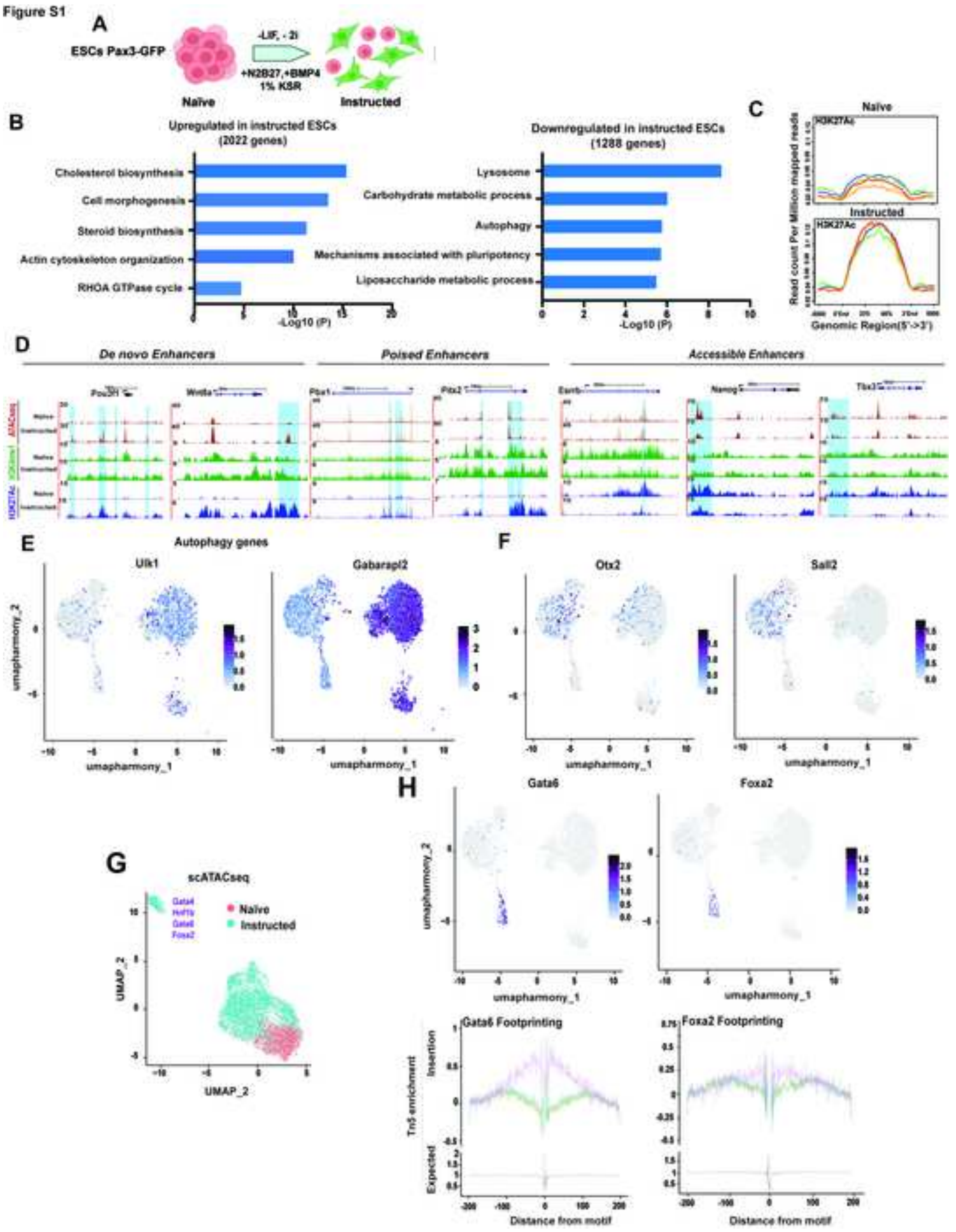

Supplemental Figure 2

Click here to access/download;Supplemental Figure;Supplemental Fig S2.tif

Figure S2

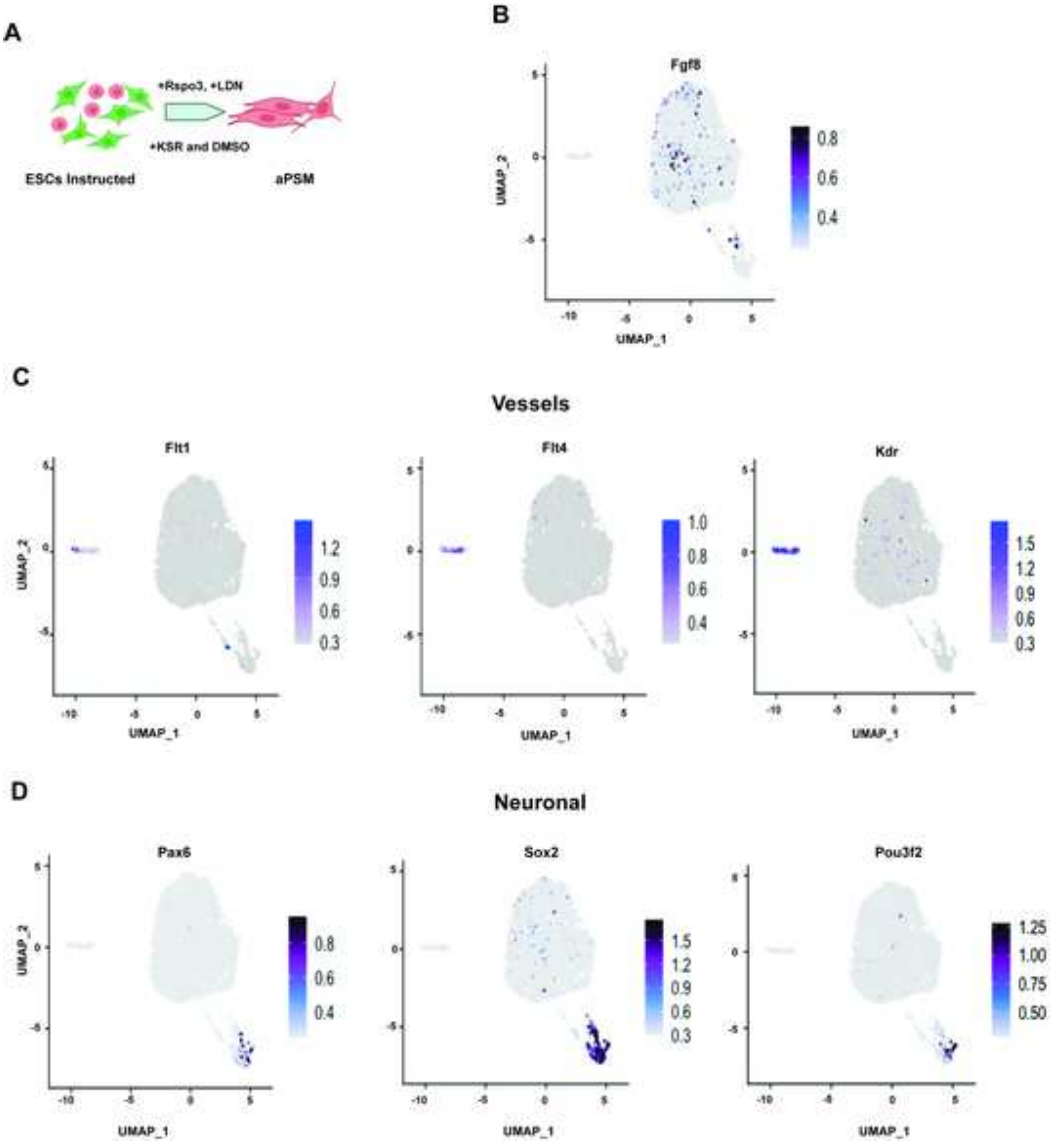

Supplemental Figure 3

Click here to access/download;Supplemental Figure;Supplemental Fig S3 .tif

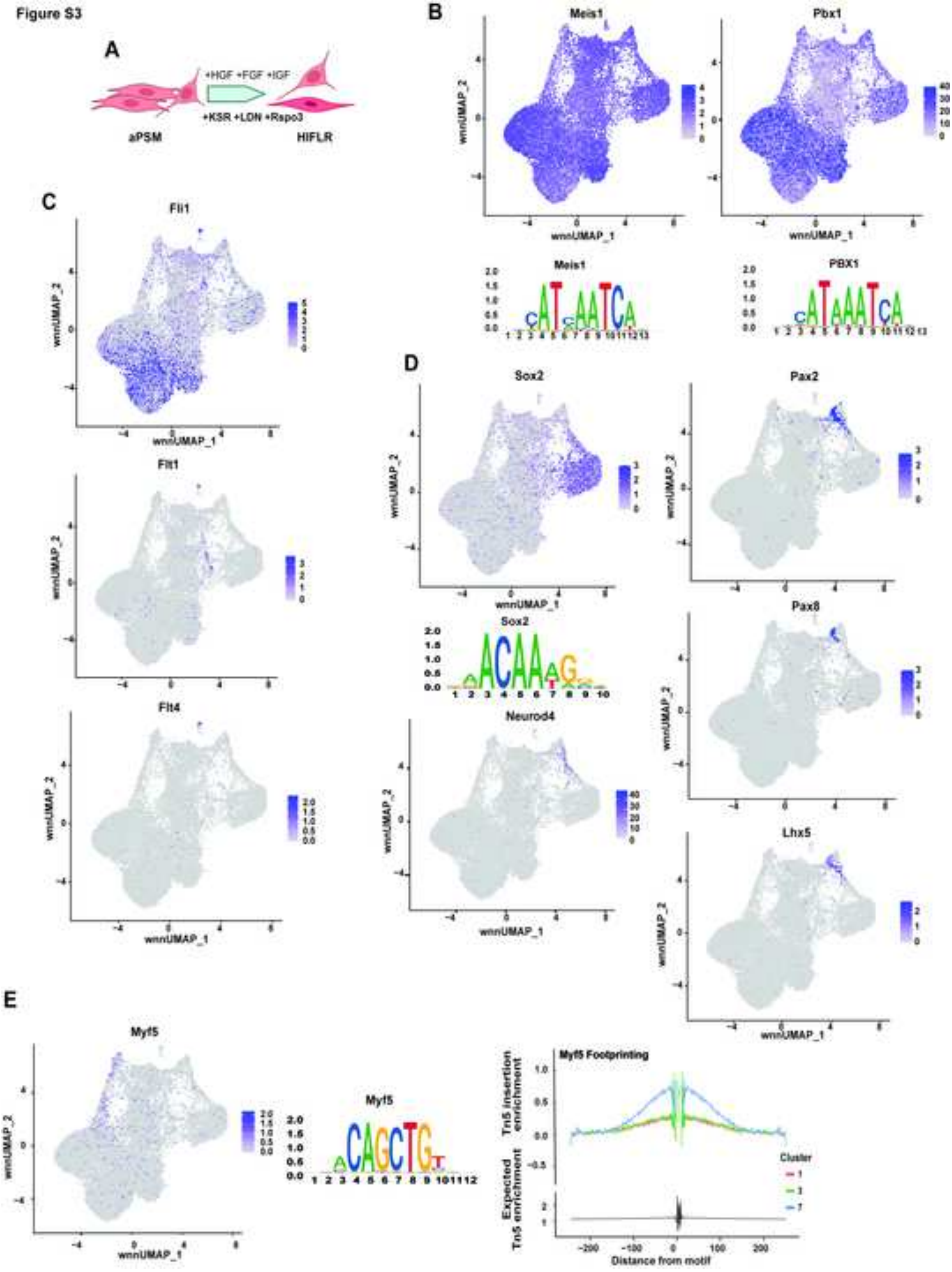

Supplemental Figure 4

Click here to access/download;Supplemental Figure;Supplemental Fig S4.tif

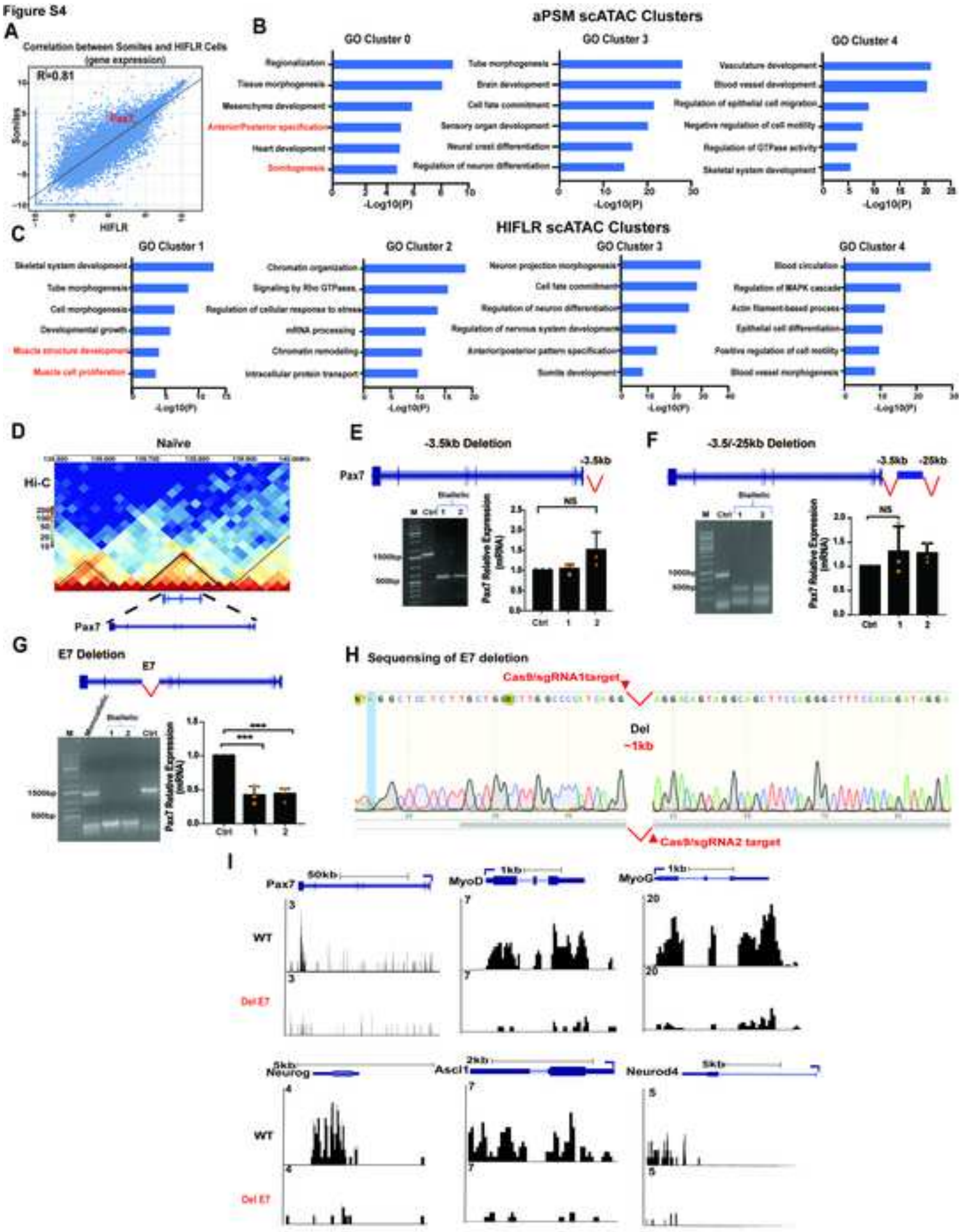

Supplemental Table S1

Click here to access/download
**Supplemental Videos and Spreadsheets**
Supplemental Table S1.xlsx

Supplemental Table S2

Click here to access/download
**Supplemental Videos and Spreadsheets**
supplemental Table S2.xlsx

Supplemental Table S3

Click here to access/download
**Supplemental Videos and Spreadsheets**
Supplemental Table S3.xlsx

Supplemental Table S4

Click here to access/download

**Supplemental Videos and Spreadsheets**

Supplemental Table S4.xlsx
